# Measuring flicker induced vasodilation at a high spatial and temporal resolution in the human retina

**DOI:** 10.1101/2024.11.20.624488

**Authors:** Pierre Senée, Léa Krafft, Inès Loukili, Daniela Castro Farias, Olivier Thouvenin, Michael Atlan, Michel Paques, Serge Meimon, Pedro Mecê

**Author notes:** These authors equally contributed to this work (Serge Meimon, Pedro Mecê).

## Abstract

Neurovascular coupling (NVC) is a crucial process in which blood flow is dynamically adjusted to meet the metabolic demands of active neurons. In this study, we introduce an innovative method for *in-vivo* imaging of retinal blood vessel dilation in response to visible light stimulation, offering insights into the functional regulation of retinal blood flow. Using high-resolution, high-speed phase contrast imaging, we continuously monitored retinal vessel dynamics in eight healthy subjects, capturing precise temporal and spatial changes in vessel size under both stimulated and baseline conditions. We have measured a significant vessel dilation of 5.2% ± 1.6% (4.4 µm ± 1.3 µm) during flicker-stimulation compared to a dilation of 2.5% ± 1% (2.1 µm ± 0.9 µm) without stimulation. The flexibility of our method also allowed for the exploration of various acquisition and stimulation settings, broadening possibilities for investigating neurovascular coupling in the retina. This work not only enhances our understanding of neurovascular coupling but also has the potential to identify new biomarkers for vision-impairing conditions and neurodegenerative diseases.

**Significance:** Neurovascular coupling is a fundamental mechanism of the brain which locally regulates blood flow in response to changes in neuronal activity. Here, we present a new imaging technique that enables measuring vasodilation in response to visible light stimulation of the neurons at high spatial and temporal resolution in the living human retina. Owing to the unprecedented measurement precision of the presented technique, we provide new insights into the inner working of functional blood flow regulation in the retina. Clinically, this method holds promise for vascular dysfunction early detection, which can improve diagnosis of retinal and neurodegenerative diseases, as well as assist the development of new targeted therapies.

## 1. Introduction

Upon activation of specific brain regions, a localized augmentation in cerebral blood flow ensues, facilitating the delivery of oxygen and essential nutrients to meet the increased metabolic demands due to neural activity. This physiological response is known as neurovascular coupling. Neurovascular coupling was first reported more than 100 years ago by Roy and Sherrington [1] and led to a large body of brain research [2], [3] where it is the source of the BOLD signal detected with fMRI [4].

Dysregulation of neurovascular coupling is observed in vision affecting diseases such as Glaucoma [5] and diabetic retinopathy [6], [7] but also in various neuropathological conditions, including ischemic stroke [8], Alzheimer’s Disease [9], and cognitive impairments stemming from hypertension [9]. This emphasizes the importance of understanding neurovascular coupling to identify new biomarkers for disease progression monitoring and to devise therapeutic strategies targeting aberrant blood flow control mechanisms following these disorders [10].

Since the retina is part of the central nervous system, neurovascular coupling can also be observed in the human retina. Administering a visible flickering light to the eye activates amacrine cells and ganglion cells in the inner retinal layers. This stimulus triggers the dilation of primary arterioles on the retinal surface, leading to enhanced blood flow throughout arteriolar, capillary, and venular networks within the retina [11]. The retina thereby provides a unique opportunity to study NVC. It shares the same embryological origin as the brain [12], [13] and as such, shares many similarities in term of structure and physiology with it. It is also the only portion of the central nervous system that is optically accessible for *in-vivo* micrometer cellular resolution imaging. Therefore, there exists the possibility that *in-vivo* retinal imaging of NVC can provide useful biomarkers of neurodegenerative disease much earlier in the disease process than conventional neuroimaging methods or clinical examination [14].

However, investigating NVC *in-vivo* remains challenging and many aspects of it are still unknown. Indeed, the dilation of blood vessels in response to light stimulation is minimal, typically only a few micrometers, necessitating a system with high resolution and contrast for accurate measurement. Furthermore, NVC is a dynamic process that requires continuous monitoring to track temporal changes in vessel diameter. Additionally, even without external stimulation, the tone of retinal arterioles is modulated by spontaneous rhythmic fluctuations in blood vessel size due to the cardiac cycle and vasomotion [15], [16], [17]. In order to be able to discriminate vasodilation provoked by flicker stimulation in the NVC process from intrinsic and spontaneous vasomotion and cardiac pulsation, an imaging method with both high spatial and temporal resolution is necessary. A better characterization of these three different sources of vasodilation is crucial not only to gain a deeper understanding of neurovascular coupling process, but also to uncover key biomarkers.

The Dynamic Vessel Analyser (DVA) (Imedos Systems UG, Jena, Germany) is the gold standard method for assessing the effect of light stimulation on retinal vessels. This system evaluates retinal vessel diameter by analyzing the brightness profile of the vessel through video sequences captured with a conventional fundus camera. It is particularly useful for measuring blood vessel dilation during a flicker test, where studies have reported a dilation of 3%-5% [18], [19], [20]. However, due to the system’s limited transverse resolution, the vessel walls are not visible, and diameter measurements rely on light absorption by the column of erythrocytes. This limitation results in reduced measurement precision, with a reported measurement sensibility of 5-7% for artery diameter, which is at the same scale as flicker induced vasodilation (Polak et al. [21]). Moreover, the DVA is only applicable to vessels with a diameter larger than 90 µm [22]. OCT technique, a gold standard in retinal imaging, can also be used to measure the diameter of blood vessels at high resolution in the axial direction. However, OCT inevitably suffer from time-varying speckle noise, which significantly impact the precision of the measurement [23], [24].

By correcting for eye aberrations, adaptive optics (AO) methods can reach transverse resolution in the retina down to 2 µm [25][26]. Owing to this resolution, blood vessel walls can be resolved [25], [26], [27], [28], [29], [30], [31], [32], [33], [34],which makes AO-based ophthalmoscopes ideal to measure the slight widening of vessels when the retina is exposed to light stimulation. Using AO-based ophthalmoscopes, researchers were able to measure the increase in lumen diameter of capillaries induced by flicker stimulation [35], [36], [37]. However, measurement of vessel caliber was only done before, during and after the flicker stimulation, thus not continuously over time. The low number of time-point measurements is due to low frame-rate, or low image contrast/ signal to noise ratio, requiring averaging for precise vessel diameter assessment. Moreover, images were acquired in a narrow field of view, making it challenging to image the same vessel or capillary over several seconds. The constraining number of time-point measurement make it not possible to account for spontaneous vasodilation process in the precise characterization of NVC.

To address these shortcomings, we developed an imaging method that enables high contrasted direct visualization of vessel wall of different calibers, arteries, and veins, at a high spatial and temporal resolution. The proposed method is the AO Rolling Slit Ophthalmoscope (AO-RSO), a method using a line scanning illumination synchronized with the rolling shutter of a 2D camera for the detection [38]. By imposing an offset between the illumination line and the rolling shutter of the camera, phase contrast images similar to those obtained with off-axis AO scanning laser ophthalmoscope (AO-SLO) [27], [28], [33], [34] can be generated. Contrary to off-axis AO-SLO, given the use of 2D camera, the AO-RSO can generate phase contrast images at high frame rate (up to 100Hz), large field-of-view (4. 5° × 2.5°) and free from motion-induced image distortion. Using the AO-RSO, we were able to reliably measure flicker induced vasodilation over time with a spatial precision of 100 nm and a temporal precision of 10-100 ms. With such high precision, we characterized the NVC response in terms of vasodilation over eight healthy subjects (**Sections 2.1 and 2.2**). Finally, we investigated how flicker-stimulated vasodilation is influenced by arterial diameter (**Section 2.3**), flicker stimulus duration (**Section 2.4**), and downstream retinal arterial areas, and venular response (**Section 2.5**).

## 2. Results and discussion

### 2.1 Measuring the Dynamics of Retinal Arteries with and without Flicker Stimulation

To measure the dynamics of retinal arteries in healthy subjects, imaging was conducted under two conditions: first, during a 40-second period with no stimulation, and second, during a flicker test. The flicker test involved 10 seconds without stimulation, 20 seconds with flicker stimulation, and another 20 seconds without stimulation. The flicker stimulation was done inside a wide rectangular 25°x20° region centered on the fovea. A green light flickering at 10 Hz and 50% duty cycle was used for all the presented studies (See **material and method** for more information). **Figure 1** presents the results for one subject and shows the example of one acquisition with **(Fig. 1G,H)** and without **(Fig. 1D,E)** flicker. An artery perfusing the fovea was chosen for imaging **(Fig. 1A)**. This was done because the region near the fovea contains a high concentration of ganglion cells [39], therefore a strong response should be observed for an artery directly perfusing this region if exposed to flicker stimulation.

**Figure 1:**
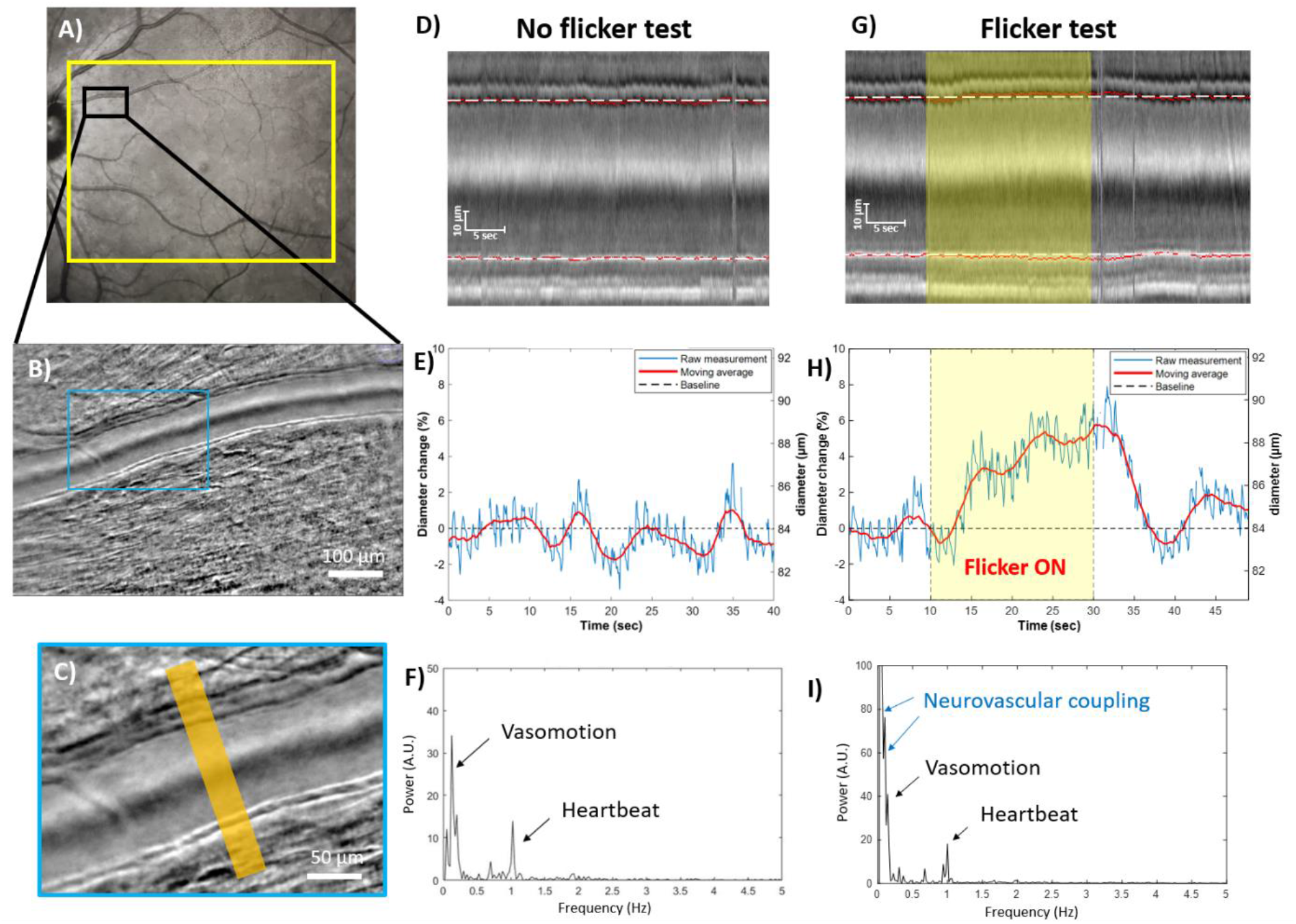
**A)** Subject’s fundus image acquired with Spectralis (Heildelberg,Germany), where the yellow rectangle indicates the stimulated area during the wide field flicker test and the black rectangle shows the imaged area. **B)** AO-RSO phase contrast image with 2.5°x4.5° FOV and high-contrasted vessel wall. **C)** Magnified region of the phase contrast image (blue rectangle), where the orange area indicates the section of the vessel where the lumen diameter is measured. **D)** M scan of the artery without flicker stimulation representing the profile of the vessel over time, white dashed lines represent the baseline position of the wall (tunica intima) of the vessel. **E)** Artery’s lumen diameter over time without flicker stimulation. **F)** Power spectral density of the lumen diameter data, where two peaks at 0.1 Hz and 1 Hz can be observed. **G)** M scan of the artery with flicker stimulation, the stimulation is represented on the M scan as a yellow rectangle. **H)** Artery’s lumen diameter over time with flicker stimulation. **I)** Power spectral density of the lumen diameter data during flicker stimulation, where additional peaks at lower frequencies (<0.1Hz) can be observed.

From the recorded images a profile of a line perpendicular to the vessel over time (M scan), representing the evolution of the vessel in time of the vessel can be constructed (**Fig. 1D,G**) and the vessel’s lumen diameter can be measured in time (**Fig. 1E,H**). The baseline diameter is defined in each acquisition as the average lumen diameter of the vessel during the first 10 seconds without flicker. Without stimulation (**Fig. 1E**), the vessel’s diameter is modulated by two components. First the cardiac cycle, responsible for the rhythmic dilation at approximately 1 Hz can be seen in blue in (**Fig. 1E,F**). Secondly, a slower oscillation at around 0.1 Hz can also be observed by looking at the moving average in red (**Fig. 1E,F**). We believe this second component to be vasomotion which is described as a spontaneous rhythmic change in the vessel diameter arising from alternating smooth muscle dilation and constriction and has been previously observed *in-vivo* [15], [16], [17]. From our study, we have seen large differences between subjects for the amplitude of vasomotion, varying from 1% to 4.8% (see supplementary information for example of vasomotion without flicker stimulation for each of eight subjects).

**Figure 1G and H** show the evolution of the diameter of an artery during a flicker test. When the stimulation starts (**Fig. 1H** at t = 10 seconds) the diameter of the artery increases rapidly at around 1% of dilation per second for approximately 5 seconds. Then the artery continues to dilate steadily at 0.1%-0.2% of dilation per second until a maximum dilation is reached around the end of the 20 seconds of flicker stimulation. After the end of the stimulation, the artery contracts back to its diameter pre-flicker. On some acquisition, this contraction undershoots under the baseline diameter before dilating back to the baseline diameter (see **supplementary information**). When looking at the power spectrum density, one can notice an additional low frequency content (<0.1Hz) compared to the spectrum without flicker stimulus, which is related to the neurovascular coupling. The general trend of the vessel’s diameter evolution during a 20 second flicker test is consistent with what has been described with fundus camera imaging using the DVA [18], [20]. However, owing to our high spatial resolution and high contrasted vessel wall visualization, our approach can, for the first time, precisely distinguish in space/time and frequency domain, the contribution of heartbeat, vasomotion and neurovascular coupling with high precision and on a single acquisition.

### 2.2 Study of the Vascular Diameter response to Flicker Stimulus on 8 Healthy Subjects

The same protocol as described in the previous section was conducted on 8 healthy subjects. For each subject, 3 acquisitions without stimulation and 3 acquisitions with a flicker stimulation were taken. For each acquisition with and without stimulation, the maximum dilation was measured. For each flicker acquisition, temporal parameters of the flicker induced response were measured as represented in **Fig. 2A**. The average and standard deviation of these measurements can be found in the table at **Fig. 2B**.

**Figure 2:**
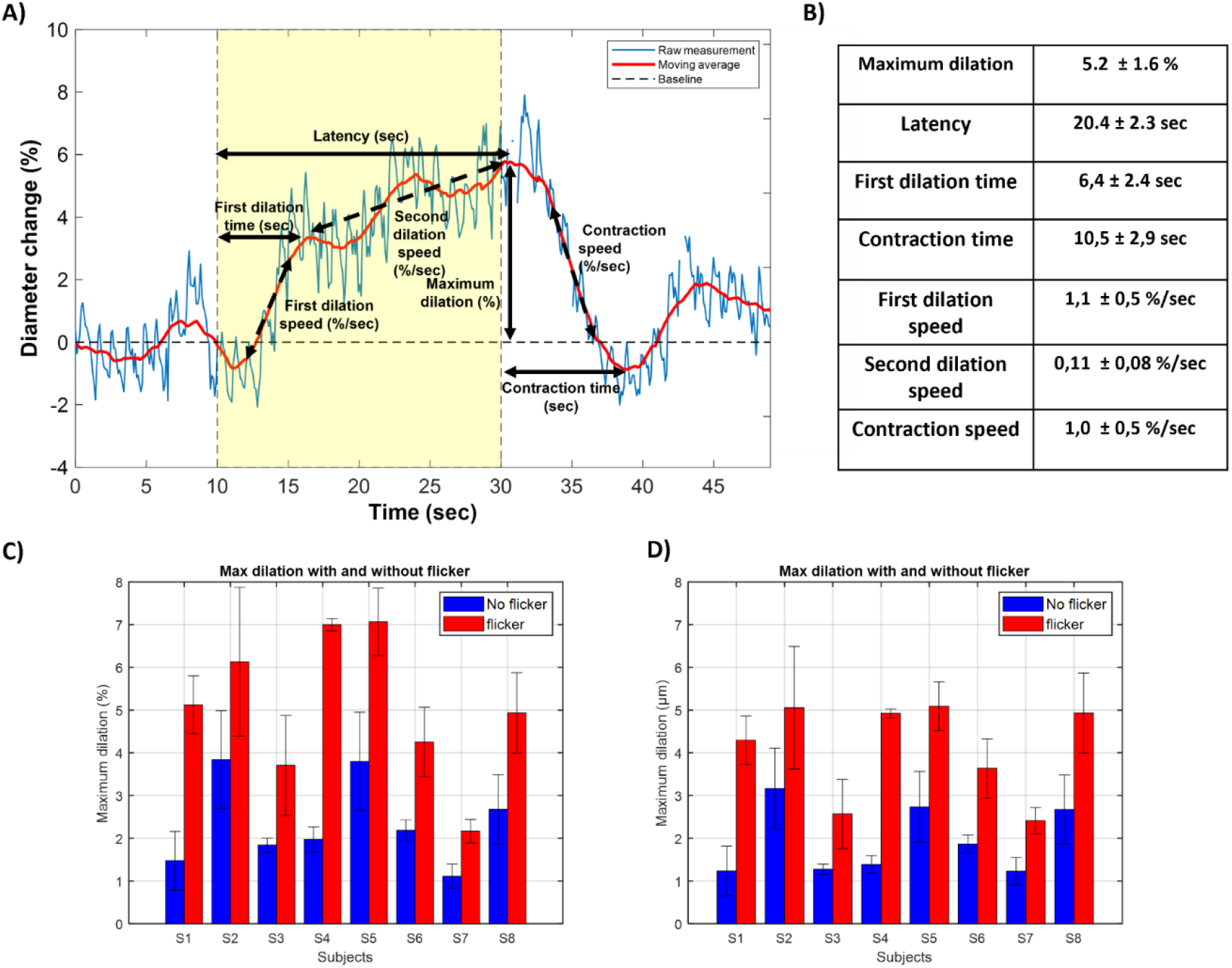
**A)** Representation of a typical flicker acquisition. Various temporal aspects of the flicker response represented on this graph, were measured for each acquisition across 8 subjects. **B)** Average and standard deviation over the population of these parameters across all acquisitions. “Maximum dilation” is defined as the peak relative increase in the lumen diameter’s moving average. The “latency” is the duration from the onset of flicker stimulation to the point of maximum dilation. “First dilation time” is the time from stimulation onset to the first maximum of the artery diameter. The “contraction time” is the interval between the end of flicker stimulation and the first minimum of artery diameter following stimulation. For each acquisition, “first dilation speed”, “second dilation speed”, and “contraction speed” were also measured in % per second. **C)** Average and standard deviation of the maximum relative (%) dilation for each subject without flicker stimulation in blue and with flicker stimulation in red. **D)** Average and standard deviation of the maximum absolute (µm) dilation for each subject without flicker stimulation in blue and with flicker stimulation in red.

When the retina is exposed to a flicker stimulation, the artery first dilates quickly at an average speed of 1.1 ± 0.5 % per second until a first maximum is reached, 6.4 ± 2.4 seconds after the onset of stimulation. Then the artery continues to dilate at a rate of 0.11 ± 0.08 % per second until a maximum dilation of the artery is attained. After the end of the stimulation the artery contracts back into its original diameter at a speed of 1.0 ± 0.5 % per second. The latency between the beginning of the flicker stimulation and the maximum dilation was of 20.4 ± 2.3 sec. On average, the maximum dilation was reached around the end of the 20 seconds of stimulation, which is consistent with previous work from Riva et al. [40] showing a saturation in the increase in blood flow in the optical nerve head after 20 seconds of stimulation.

**Figs. 2C and D** represent the relative and absolute maximum dilation on all the acquisitions on the 8 subjects respectively. For all the subjects, the maximum dilation was significantly higher with the flicker stimulation (red bars in **Fig. 2C and D**) than without the flicker stimulation (blue bars in **Fig. 2C and D**). A 5.2 % ± 1.6 % (4.4 µm ± 1.3 µm) maximum dilation was recorded during the flicker acquisitions on the 8 subjects. The maximum dilation without flicker stimulation was of 2.5 % ± 1% (2.1 µm ± 0.9 µm). We believe this observed dilation without stimulation to be due to vasomotion. **Figures 2C and D** also show a variability of maximum dilatation with flicker stimulation among subjects (varying from 2.2% to 7.1%) as well as a variability among different acquisitions for a single subject (varying from 0.3% to 1.7%). The intra- and inter-subject variability can be attributed to the occurrence of vasomotion and its large amplitude variability, as well as intrinsic different NVC response among subjects. Another possible explanation is the artery caliber, as some studies suggest that smaller arteries and arterioles have a more significant vessel dilation in response to flicker stimulation [41]. Given the spatial selectivity of neurovascular coupling in the retina [35], the composition (containing more or less activated neurons) of the region which is perfused by the artery may also influence the vasodilation response. To get a better understanding of the factors which can determine the strength of this response we investigated in later sections the effect of the size of the artery (**Section 2.3**) and the effect of the region perfused by the artery (**Section 2.5**).

### 2.3 Influence of Arterial lumen Diameter on Flicker Induced Vasodilation

We investigated the influence of the diameter of the artery on the response to flicker stimulation. In **Fig. 3A** are represented the maximum dilation as a function of baseline arterial diameter. A clear trend could be seen where the smaller the artery, the larger the dilation, with a coefficient of determination *R*^2^ = 0.54, showing a linear correlation. To get more insight into the temporal aspect of the dilation depending on the size of the arteries, we separated the subjects into three groups depending on the baseline diameter of the imaged arteries. In **Fig. 3B** the average curves from the three groups are plotted. Notably, the maximum dilation is most pronounced in the d < 80 µm group and least pronounced in the d > 90 µm group. Moreover, directly after the onset of flicker, in the initial fast dilation phase, the three groups exhibit a drastically different dilation speed. As illustrated in Fig. 3B, the d < 80 µm group dilates at a rate of approximately 1% (0.7 µm) per second, the 80 < d < 90 µm group dilates at a rate of approximately 0.7% (0.55 µm) per second, and the 90 µm < d group dilates at a rate of 0.4% (0.4 µm) per second.

**Figure 3:**
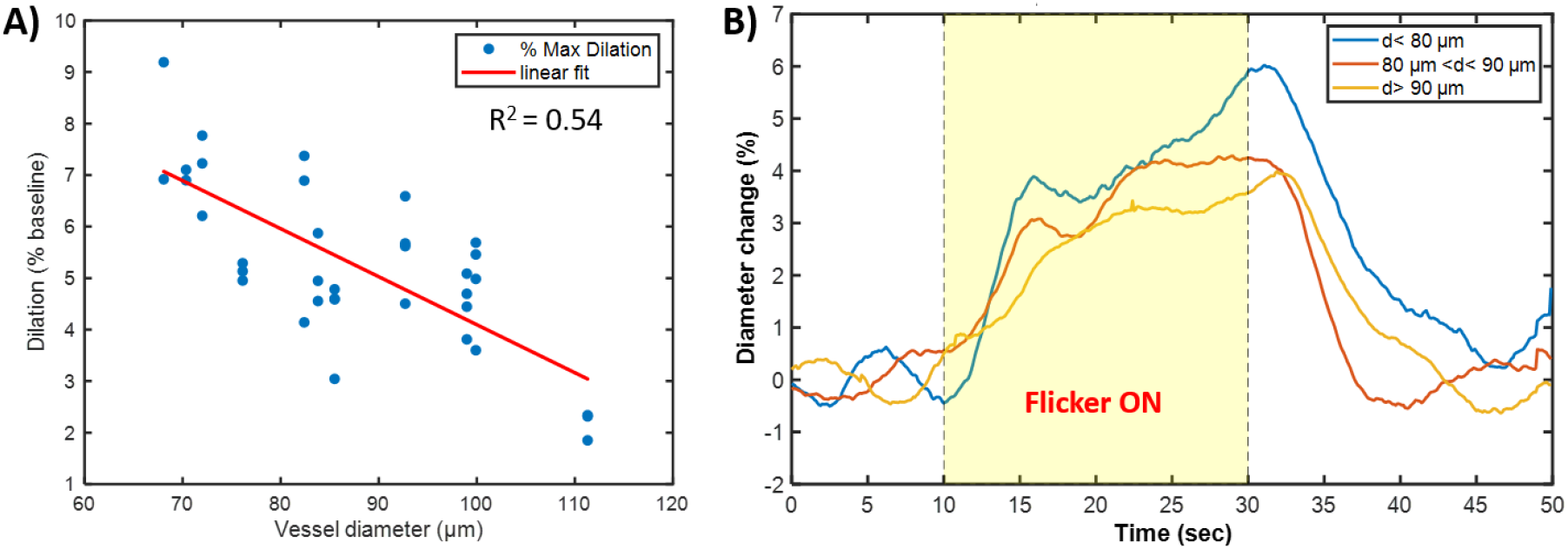
**A)** Maximum dilation for each flicker stimulation as a function of baseline lumen diameter. **B)** Averaged graph of every flicker test grouped by baseline lumen diameter “d”: blue: < 80 µm, red: between 80µm and 90µm, yellow: >90µm.

### 2.4 Studying the Effect of Flicker Duration

We used our system to study the effect of the duration of the flicker stimulation on flicker induced vasodilation. On the same subject, we conducted flicker test of increasingly longer stimulation times (0,2,5,10,20,40 and 60 seconds). For each acquisition, the system imaged an artery upstream of the fovea for 10 seconds without stimulation followed by the flicker stimulation, followed by 20 seconds without stimulation. Three acquisitions were taken for each flicker stimulation duration and then averaged. The results from these acquisitions are presented in **Fig. 4A**.

**Figure 4:**
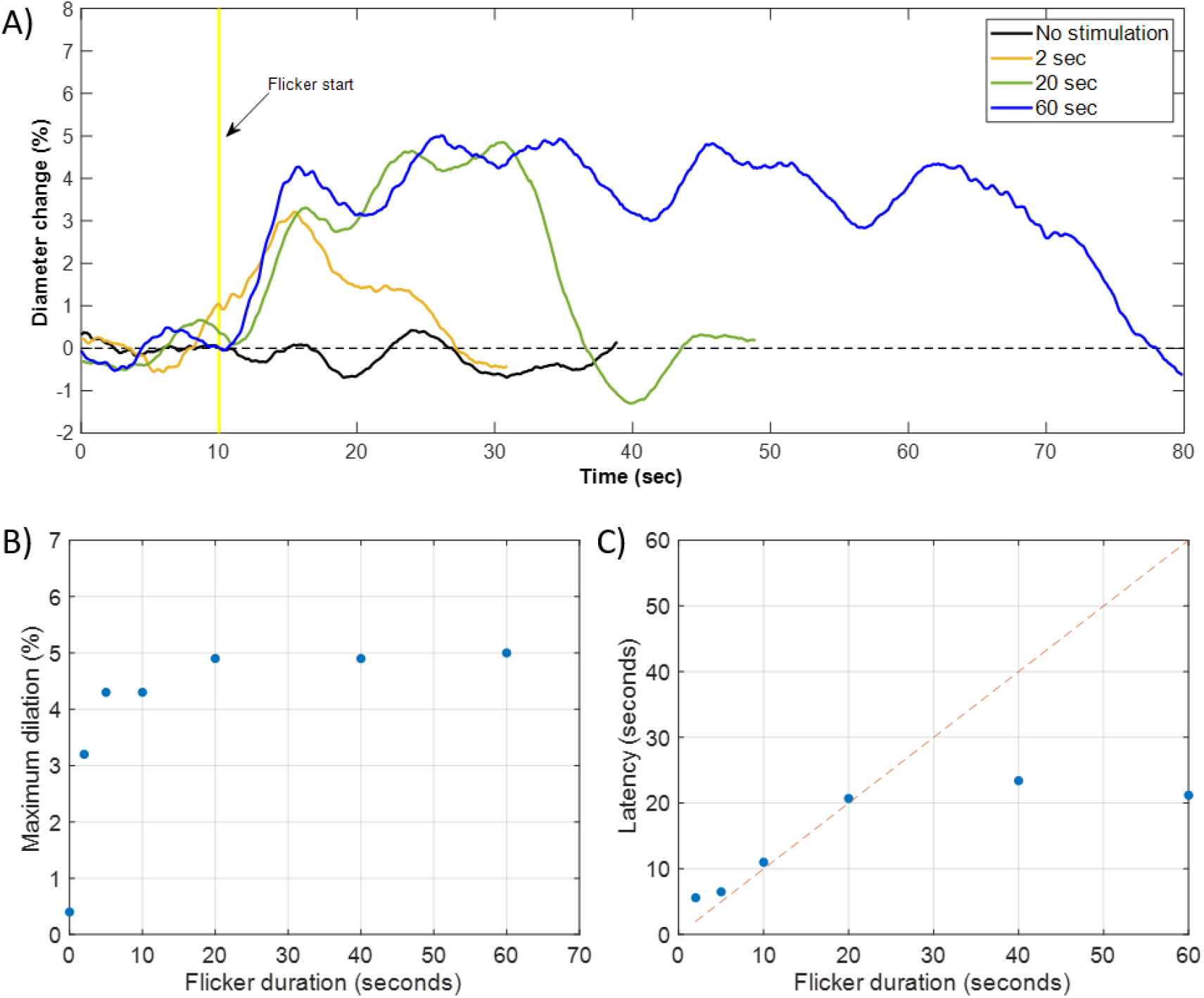
**A)** Moving average lumen diameter curves for different flicker durations, with flicker stimulation starting at t = 10 seconds. For clarity, only flicker durations of 0, 2, 20, and 60 seconds are represented. **B)** Average maximum dilation across all flicker duration conditions. **C)** Average latency of dilation across all flicker duration conditions, the dashed representing a linear behavior.

For very short stimulation durations, although a significant increase in the vessel’s diameter can be observed compared to without stimulation, the dilation is less pronounced and with a different shape compared to the one with the standard 20 seconds of flicker stimulation. As shown in **Fig. 4A**, a ∼3% maximum dilation for 2 seconds of stimulation was obtained. Interestingly, the maximum dilation is reached around 6s after the beginning of the flicker stimulation, therefore 4s after the end of the flicker. This delayed dilation is probably linked to the first dilation observed for 20s flicker duration described in **Fig. 2A**. Such phenomenon may indicate that the first dilation and the second dilation have different origins. The first fast dilation seems to be independently of the light stimulation duration, with a peak around 6s after the beginning of the stimulation. On the other hand, the second slower dilation only happens if the duration is longer than 6s.

For very long stimulations, the artery does not dilate more than the maximum dilation reached for 20 seconds of stimulation, as can be seen in **Fig. 4B**. For 40 and 60 seconds of stimulation, the vessel’s diameter reaches a plateau after ∼20 seconds, which is reflected in the latency measured in **Fig. 4C**. For some of these long stimulations, the vessel’s diameter starts to slightly decrease even before the end of the stimulation as it can be observed for 60 seconds duration in Fig. 4A.

This preliminary study shows that 20 seconds duration flicker stimulation is optimal to achieve maximum vasodilation. Interestingly, we also highlighted that very brief stimulations of just a few seconds can produce significant vasodilation compared to the baseline, where only the first fast dilation (as described in **Fig 2A**) is visible. A short flicker test to assess NVC function could benefit clinical examination, as it could be to more comfortable for the patient for a faster examination and image processing.

### 2.5 Study of simultaneous flicker response of one vein and two arteries perfusing different areas of the retina

We took advantage of our system’s large FOV to measure the effect of flicker stimulation on three vessels simultaneously. On one subject, we selected an area of interest where two arteries and one vein could be imaged within a 4.5°x4.5° FOV. This area was chosen because it contains two large arteries and one large vein. Additionally, the two arteries supply blood to distinct regions. As illustrated in Fig. 5A, artery 1 directly supplies the fovea, whereas artery 3 supplies the peripheral retina. The diameter of all three vessels was measured and their evolution over time are represented in **Fig. 5**.

**Figure 5:**
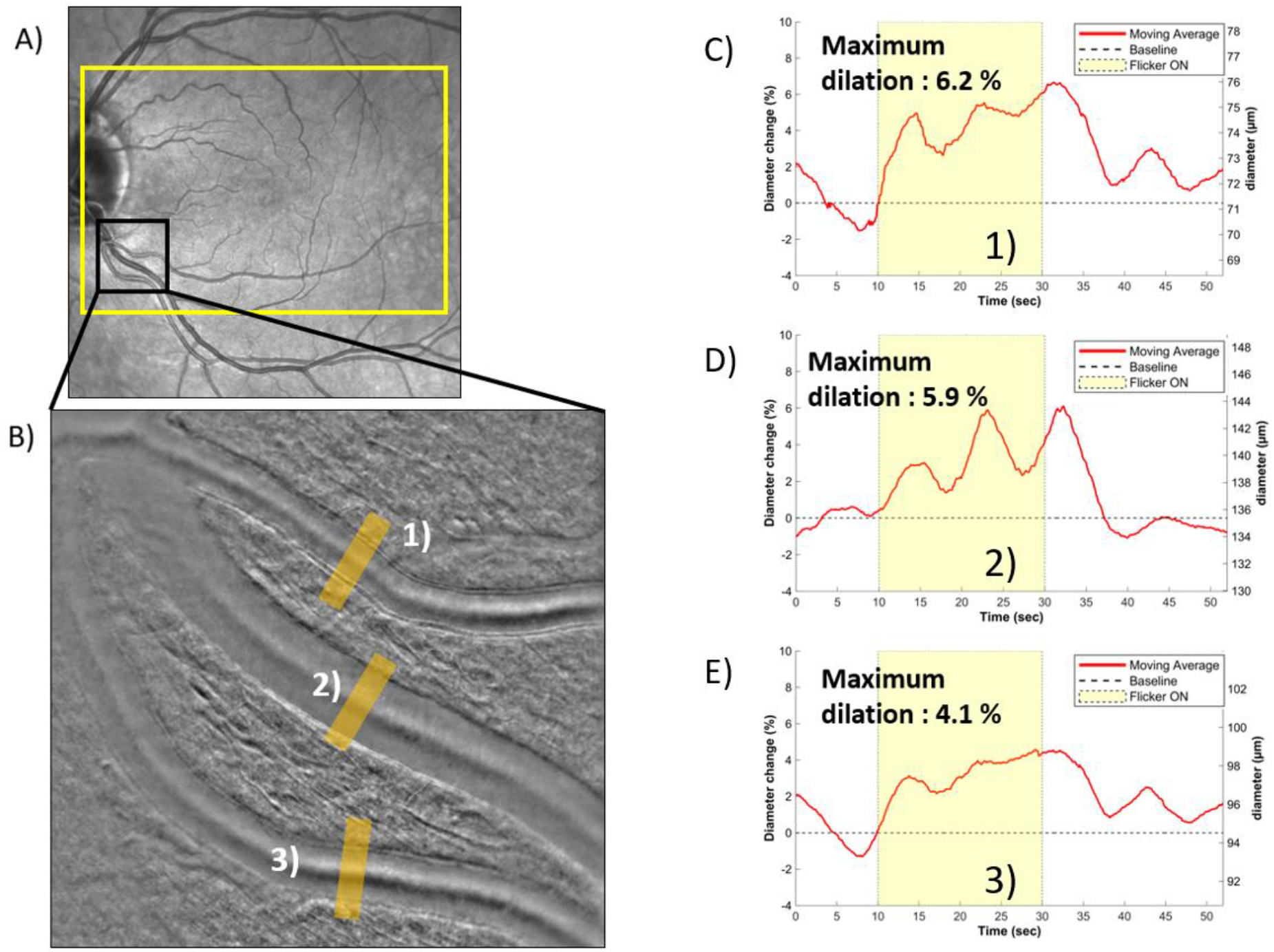
**A)** Subject’s fundus image acquired with Spectralis (Heildelberg,Germany), where the yellow rectangle indicates the stimulated area during the wide field flicker test and the black rectangle shows the imaged area. **B)** AO-RSO phase contrast image, with a FOV of 4.5°x4.5°, where we can identify three vessels: two arteries (1 and 3) and a vein (2). The area where the lumen diameter was measured is represented with an orange rectangle. **(C**,**D**,**E)** Evolution in time of the lumen diameter of vessels 1,2 and 3 respectively.

We can observe that the vasomotion between the two arteries in **Fig. 5C,E** is mostly synchronized. This resemblance may come from their shared origin as daughter branches of the same artery. However, artery 1, which perfuses directly the fovea, exhibits a notable dilation of 6.2%, while artery 3, supplying the periphery, dilates only up to a maximum of 4.2%. With the ganglion cell layer being thicker near the fovea than it is at the periphery of the retina, this observation is in line with the known phenomenon of neurovascular coupling enhancing blood flow specifically to the activated regions in response to flicker stimulation [35]. However, artery 1 has also a smaller baseline diameter than artery 3. As we have seen in the previous section, small arteries tend to dilate more than larger ones. More investigation needs to be done to understand which effect is predominant. The temporal evolution of the vein lumen diameter differs significantly from that of the two arteries **(Fig. 5D)**. The increase in diameter following stimulation appears more gradual and exhibits larger vasomotion fluctuations compared to the arteries. Additionally, a slight delay in the dilation of veins compared to arteries is observed, which aligns with the arteries’ more active role in blood flow regulation, in contrast to the veins’ more passive role.

## 3. Conclusion

We developed a high resolution, high frame rate, large field of view phase contrast imaging system to characterize with unprecedented spatiotemporal precision, the effect of flicker light stimulation on large arteries and veins of healthy subjects. This system was employed to assess the effects of flicker stimulation on the diameter of large arteries and veins (ranging from 67µm to 110µm), in a cohort of eight subjects. The proposed method enabled reliable measurement of flicker-induced vasodilation with a spatial resolution of 100 nm and a temporal resolution of 100 ms. Additionally, the large field of view and distortion-free imaging provided by the system ensures high robustness, enabling continuous data acquisition for up to four minutes, with data loss occurring exclusively during subjects’ blinks and micro-saccades.

The complete temporal dynamics of retinal artery diameters were measured in eight healthy subjects, both with and without flicker stimulation. Our imaging system allowed simultaneous observation of the effects of the cardiac cycle, vasomotion, and flicker stimulation on the measured vessel diameter. During flicker stimulation, a significant maximum dilation of 5.2% ± 1.6% (4.4 µm ± 1.3 µm) was observed across all subjects, compared to a maximum dilation of 2.5% ± 1% (2.1 µm ± 0.9 µm) without stimulation. Continuous high-speed measurements revealed the modulation of retinal vessel diameter by both the cardiac cycle and vasomotion, emphasizing the necessity of capturing the full temporal profile of dilation to study neurovascular coupling effectively, rather than relying solely on pre- and post-stimulation diameter measurements, as it is commonly done in retinal neurovascular studies.

Furthermore, various characteristics of the flicker-induced vasodilation response were investigated. Across subjects, a correlation was observed between flicker-induced vasodilation and arterial size, with smaller arteries demonstrating greater maximum dilation compared to larger arteries. The effect of flicker stimulation duration was also evaluated in one subject. For short stimulation durations (<20 seconds), a significant increase in vessel’s diameter was noted, although the response was less pronounced compared to the standard 20-second stimulation. Conversely, for stimulation durations longer than 20 seconds, no further dilation was observed beyond the maximum achieved with 20 seconds of stimulation.

The system’s large field of view was also used to image two large arteries and one large vein in a single subject, allowing for the simultaneous assessment of vasomotion and flicker-induced vasodilation in vessels perfusing different retinal regions. Interestingly, of the two arteries examined, the one perfusing the fovea exhibited a greater maximum dilation compared to the artery supplying the retinal periphery, suggesting as expected that neurovascular coupling preferentially enhances blood flow to regions of the retina with a higher density of activated neurons.

While additional acquisitions on multiple subjects would be required to establish a precise characterization of flicker induced vasodilation on different populations and clinical cases, the presented method and findings pave the way for further exploration of NVC at sub-micrometer and millisecond resolution. Indeed, a precise characterization, in space and time, of the mechanisms underlying physiological and/or pathophysiological NVC responses can facilitate current queries in neurovascular metabolism research, could provide new biomarkers for the evaluation of retinal and neurodegenerative diseases, and may ultimately lead to the development of therapies to prevent the breakdown of NVC.

## 4. Material and Method

### 4.1 Subjects

Eight subjects, ranging in age from 24 to 33 y (S1 = 24, S2 = 25, S3 = 26, S4 = 25, S5 = 27, S6 = 27, S7 = 28 and S8 = 33 years old) and free of ocular disease, participated in the experiments. All subjects had a spherical equivalent refraction between 0 and −4.5 diopters. All had normal intraocular pressure (IOP), and appearance of optic disk and fundus. Written informed consent was obtained from all participants following an explanation of experimental procedures and risks both verbally and in writing. All experiments adhered to the tenets of the Declaration of Helsinki. The total irradiance for the imaging and AO light sources was, respectively, 1700 μW and 3.8 μW, which is below the ocular safety limits stipulated by the International Organization for Standardization (ISO) standards for group 1 devices.

### 4.2 Delivery of Flicker Stimulation

A stimulation screen (Waveshare, China) was integrated into the imaging system. The screen provided a 25°x20° illumination region centered on the fovea. The screen was combined with the imaging system using a pellicle beam splitter. A programmable, stable fixation target was provided for the subject during all stimulus conditions. The flicker stimulation was generated and delivered using a custom Python software allowing the simultaneous start of the recording with the imaging system and the beginning of the flicker test protocol. The full-field flicker stimulus used in this experiment was green (530 nm, 40 nm FWHM) with a square-wave modulation of the entire field at 50 % duty cycle with a maximum illuminance of 7.3 lx and a minimum illuminance of 0 lux. These illuminance intensities and modulation rates were chosen to be similar to previous literature studying neurovascular coupling in the retina in order to induce high metabolic activity [40], [42], [43].

### 4.3 Imaging System

We used our Adaptive Optics Rolling Slit Ophthalmoscope (AO-RSO) which has been previously described elsewhere [38]. In short, it consists of a 2D camera based, non-confocal, split detection AO-line scanning ophthalmoscope. The light emitted from a SLED (Thorlabs) is projected in the retina into a line pattern of 10 µm width using a powell lens. The light backscattered by the retina is then detected using the rolling shutter of the detection camera (ORCA-Fusion, Hamamatsu). The rolling shutter and the galvanometer mirror of the illumination scan are synchronized via a custom Matlab (Mathworks, Natick, MA) software. To achieve phase contrast imaging of the vessel walls, a 10 µm offset between the illumination laser line and the exposed pixels of the rolling shutter is imposed. The exposure time of each line of pixel of the rolling shutter is chosen at 300 µs, thus giving an effective width of the rolling shutter of 50 µm in retinal space. Consecutive positive and negative offset images are alternated and subtracted to achieve split detection imaging, further increasing the phase contrast of images. The AO-RSO allows to acquire images at high resolution (2 µm), high speed (100 Hz), using phase contrast imaging on a large field of view (2.5°x4.5°), and without motion-induced image distortion. Finally, owing to the vertical scanning, one can increase the FOV by sacrificing the frame rate (e.g. 4.5°x4.5° for 50 phase contrast images per second) or a shorter FOV for a faster acquisition rate.

### 4.4 Image Acquisition Protocol

All imaging sequences were collected with the room lights off. The subject’s eye was cyclopleged and dilated using tropicamide 0.5%. The eye and head were aligned with the imaging system using a chinrest mounted to a motorized XYZ translation stage. Correct imaging focus was realized by optimizing vessel walls contrast using the real-time displayed images. For each subject, an artery upstream of the fovea in the nasal region was selected using fundus camera image of the subject. The imaging region was chosen along the selected artery and at ∼8° eccentricity. Once the imaging region was selected, three acquisitions of 40 seconds without stimulation were taken. Then three flicker tests, each consisting of 10 seconds of recording without stimulation, 20 seconds of flicker stimulation and 20 seconds without stimulation were taken. Between each flicker test, a one-minute break was included.

Recording by the imaging system was synchronized with the stimulation channel. The system acquired images continuously during the entirety of each 40-second baseline recording and during each 50-second flicker tests. The system provides phase contrast imaging at 100 Hz on a 2.5 × 4.5° FOV allowing visualization of 1.2 mm sections of horizontal blood vessels and making it possible to keep the vessel of interest inside the imaging field of view despite imperfect eye fixation which is unavoidable during 50 sec acquisitions.

### 4.5 Image Processing

Images were first registered using a custom Matlab (Mathworks, Natick, MA) phase correlation algorithm [44]. Images were averaged in groups of 10 to enhance signal-to-noise ratio, thereby producing a sharp image of the blood vessels every 100 ms. This approach was employed to enhance signal-to-noise ratio while maintaining the capability to monitor blood vessel’s fluctuation in diameter due to the cardiac cycle in all subjects.

### 4.6 Vessel Walls Detection and Measurement

From the registered and averaged images, a 30 µm segment of the vessel was chosen. For each averaged image, pixel lines perpendicular to the vessel’s direction within the selected segment were averaged. These averaged lines, each representing the cross-sectional average of the vessel in the 30 µm segment, were plotted over time to produce an M-scan of the vessel (Fig. 1C, 1E). From the M-scan, the positions of the vessel’s inner and outer wall were measured on each side. A MATLAB peak detection algorithm was used to determine their positions by identifying the two maxima and two minima in each pixel line of the M-scan corresponding to the tunica intima and the tunica externa of both sides of the vessel. The lumen diameter was defined as the distance between the two inner walls of the vessel. A parabolic fit was implemented to achieve a subpixel measurement of the lumen diameter with a precision of 0.1 µm. To represent the measured diameter, both the raw measurements taken every 0.1 seconds and the moving average were represented. The moving average was chosen on a 3 second window to filter out the effect of the cardiac cycle. For the measurement of the maximum dilation (represented in Fig 2A), the moving average was chosen instead of the raw measurement, this was done to filter out the effect of the cardiac cycle on the lumen diameter which variations don’t reflect the effect of neurovascular coupling on the vessel.

## Funding

Office National d’études et de Recherches Aérospatiales (PRF TELEMAC); Agence Nationale de la Recherche (ANR-18-IAHU-0001,ANR-22-CE19-0010-01).

## 6. Supplementary Information

Here, we present one example of acquisition with and without flicker stimulation for each of the eight healthy subjects imaged in this study (S1-S8). In all presented supplementary figures, the reader will find their fundus photography image (A), where the yellow rectangle represents the stimulated area during the wide field flicker test and the black rectangle represents the imaged area, which is highlighted below as a magnified image (B). The profile of the vessel over time without flicker stimulation can be observed in (C), where the dashed white lines represent the baseline position of the wall (tunica intima) of the vessel. (D) shows the artery’s diameter over time without flicker stimulation. The vessel profile and vessel diameter over time during flicker stimulation can be seen in (E) and (F) respectively.

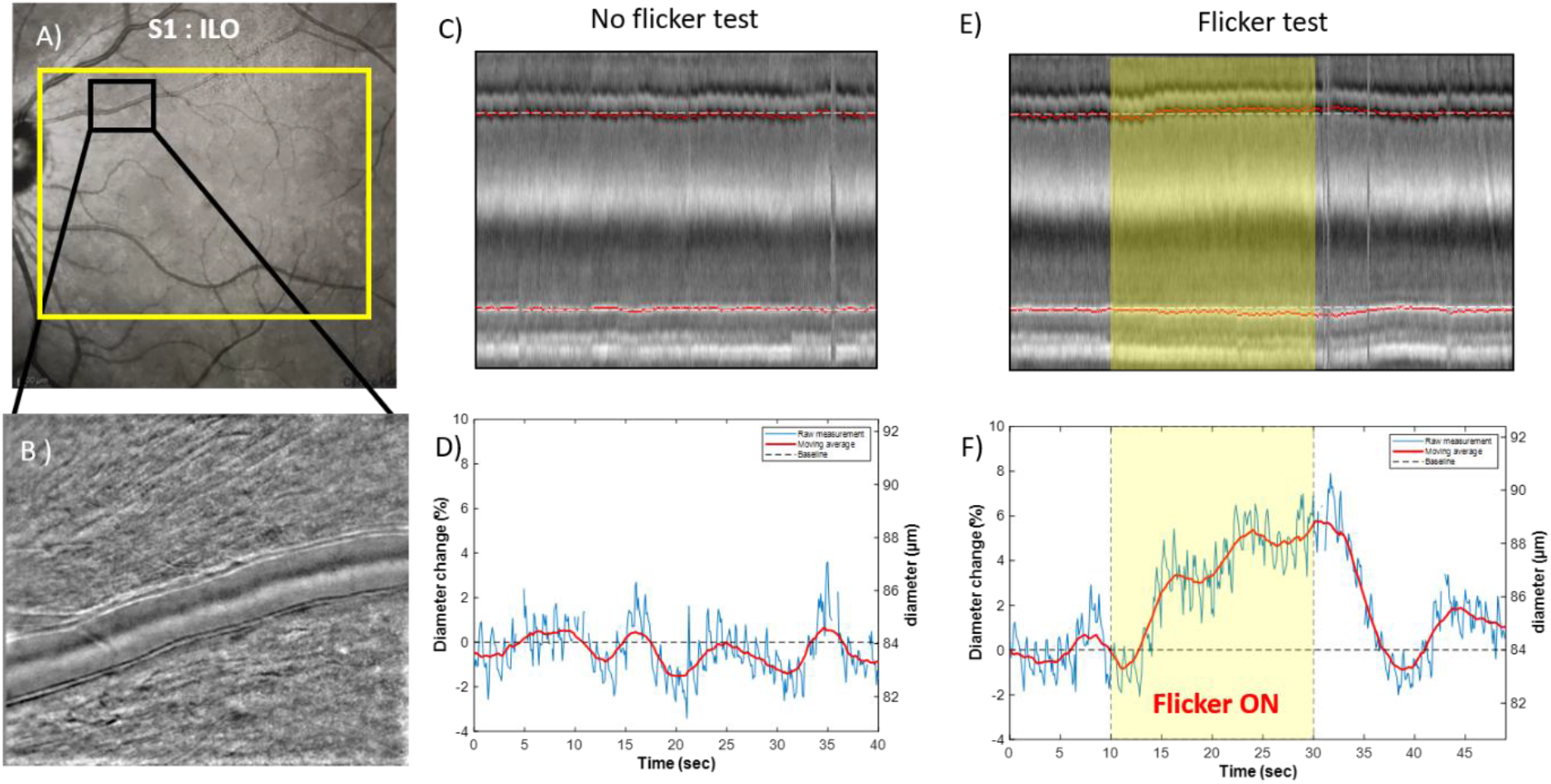

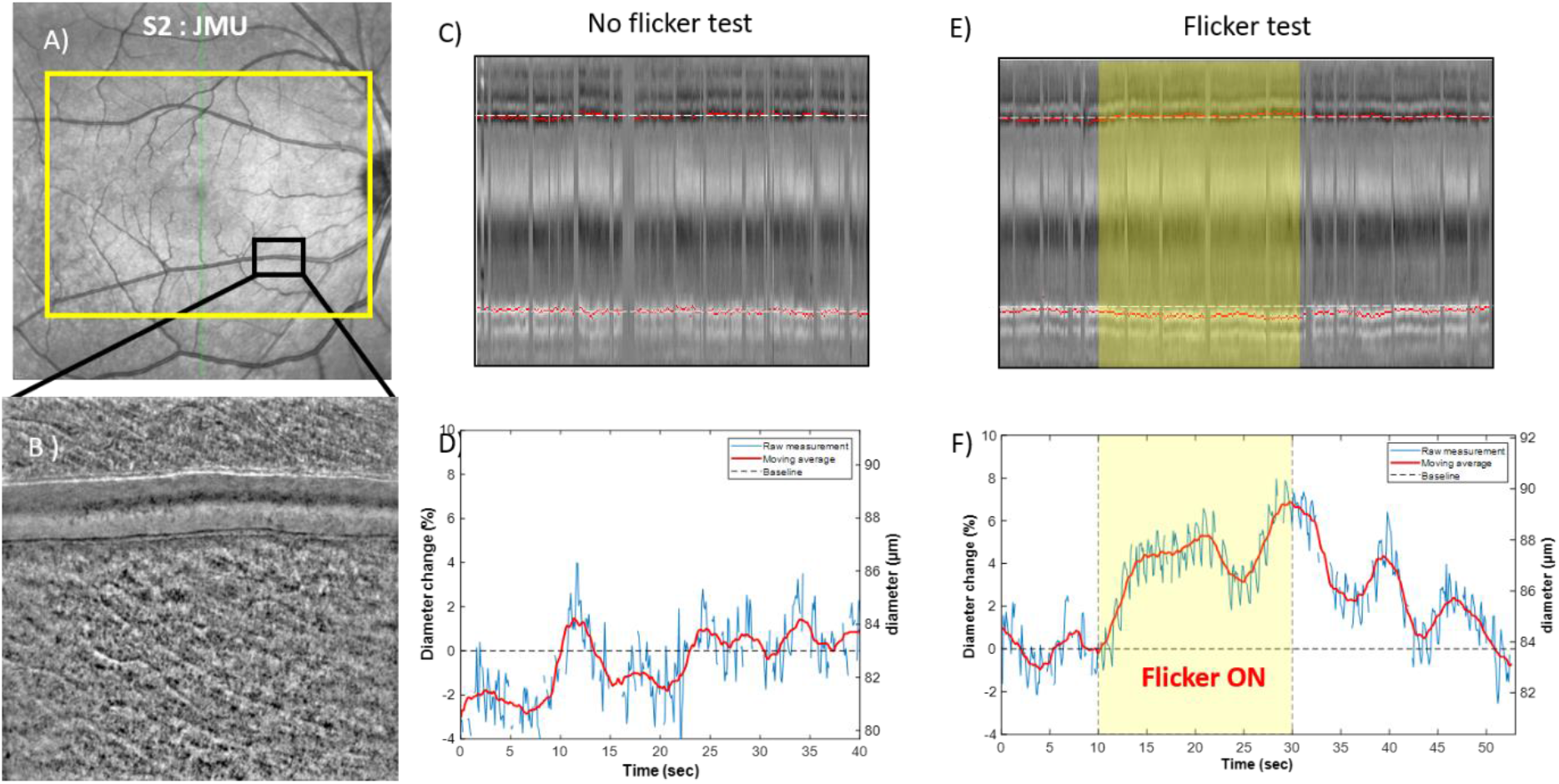

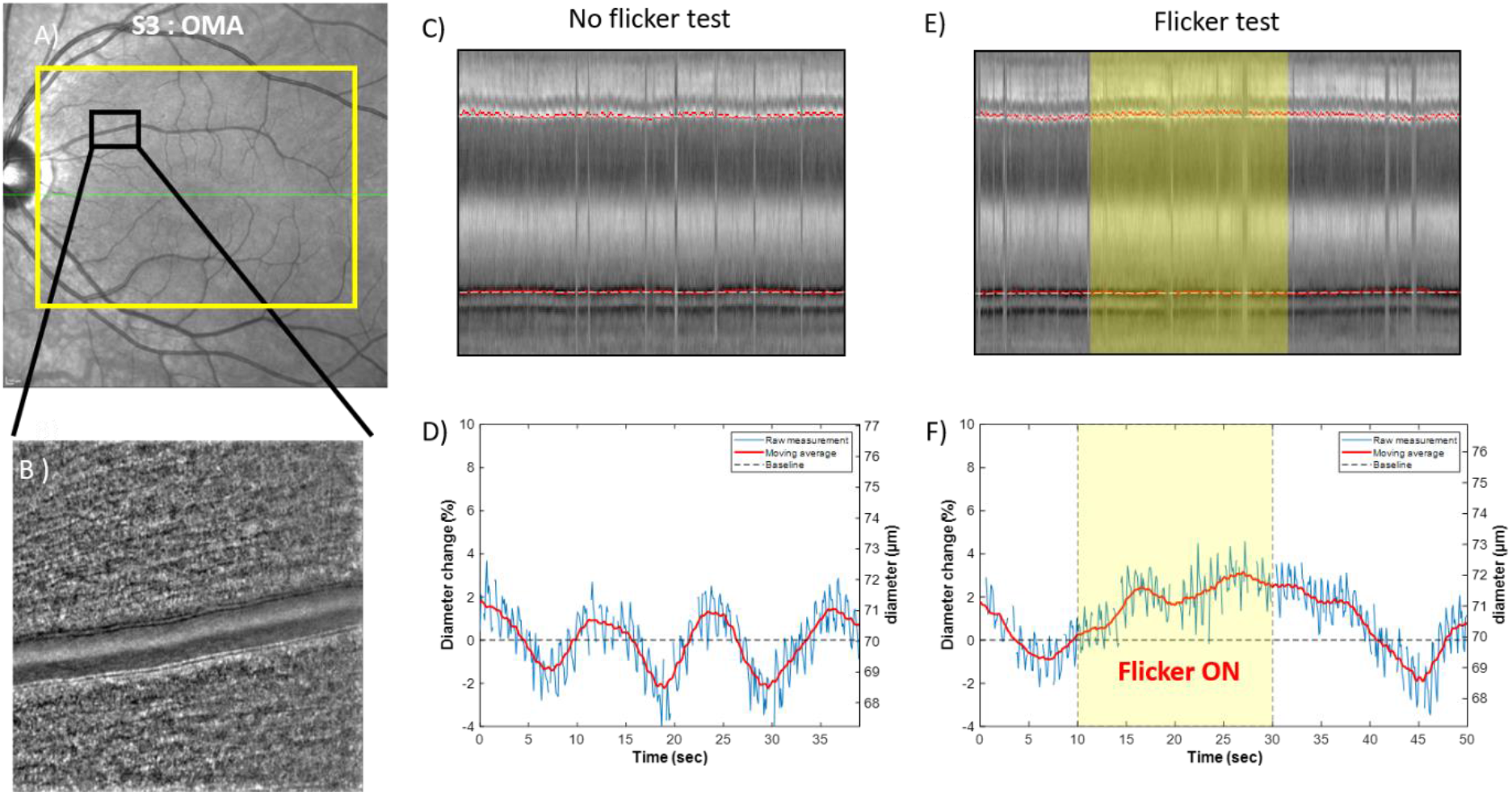

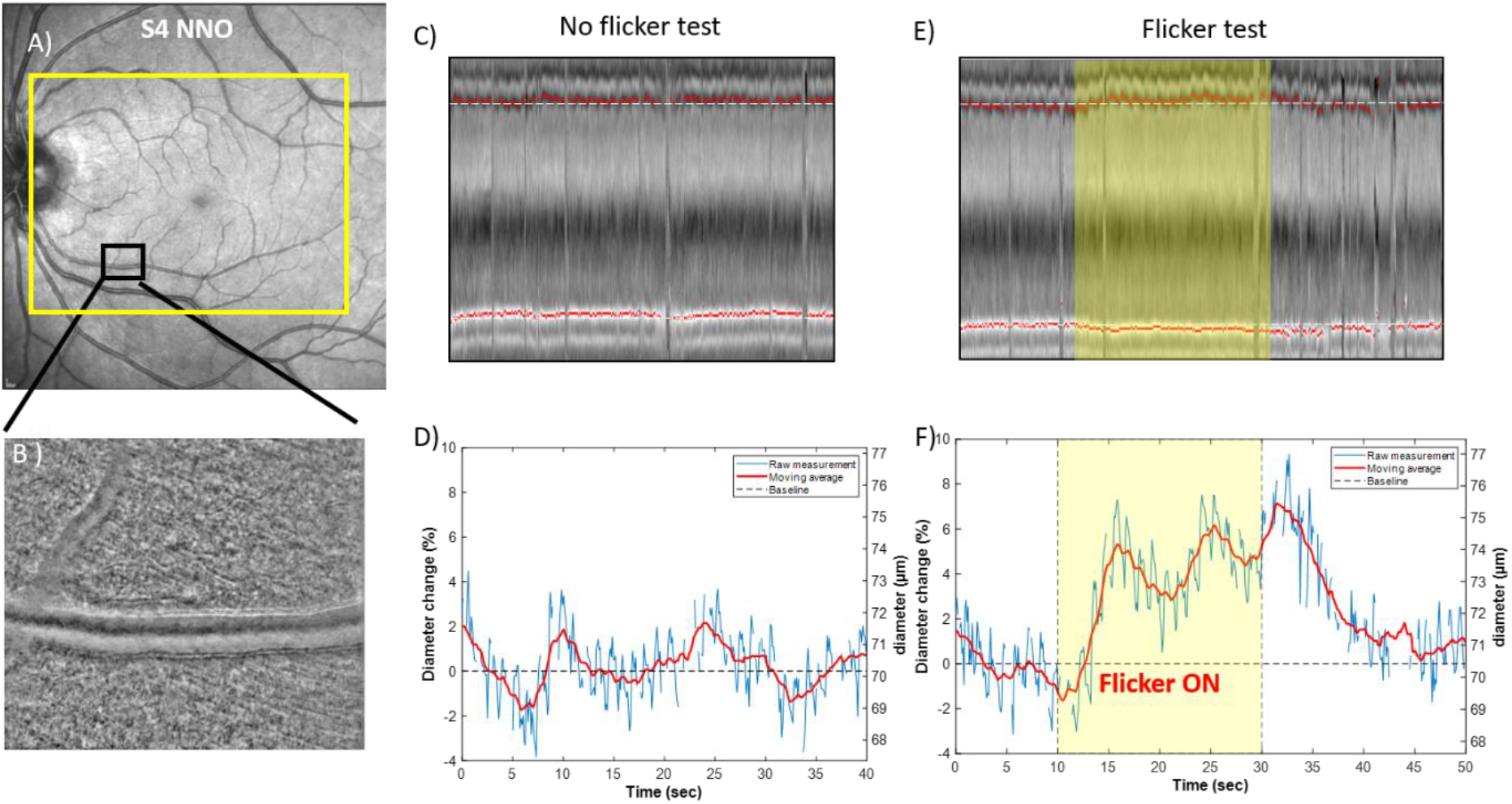

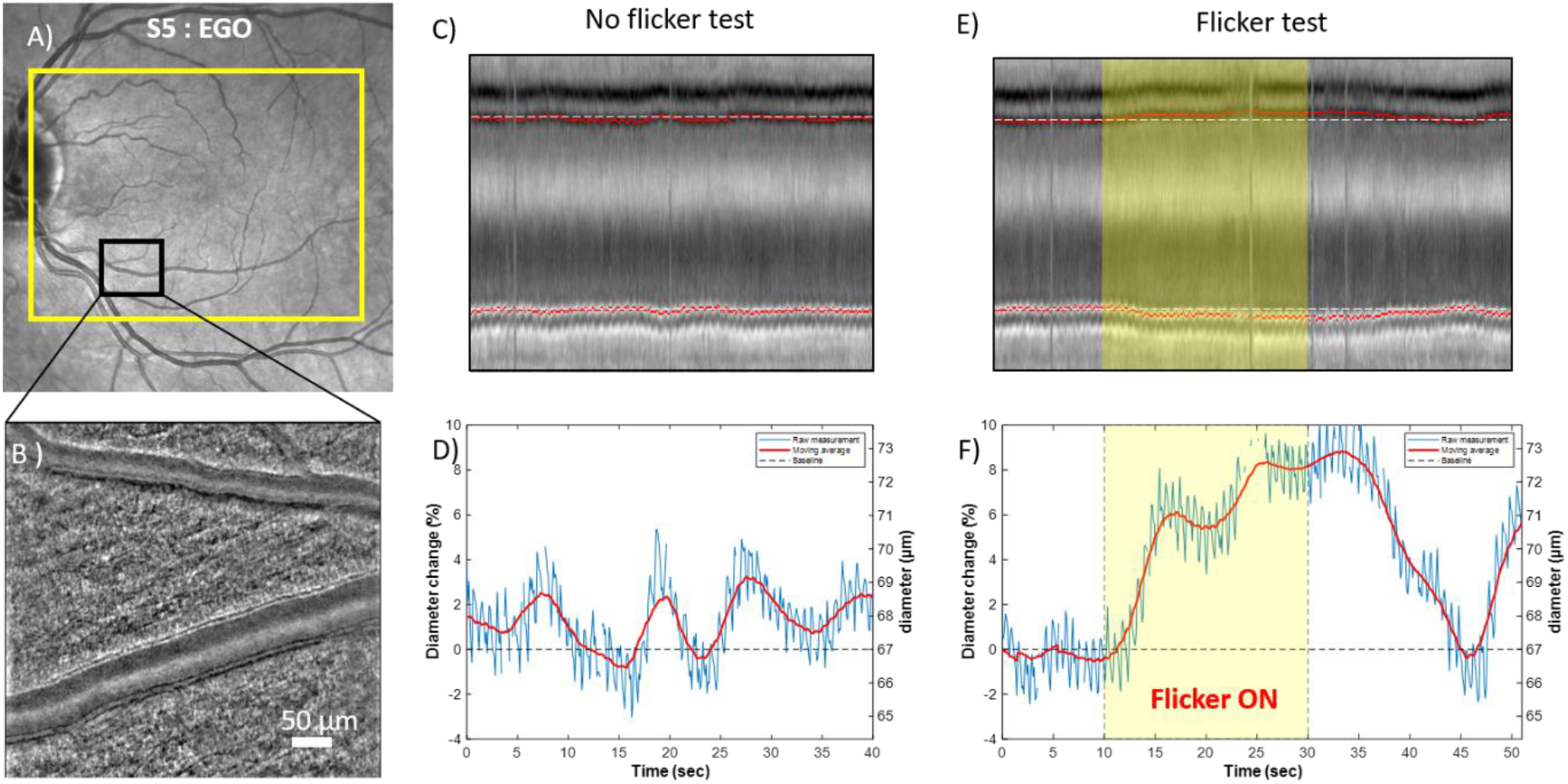

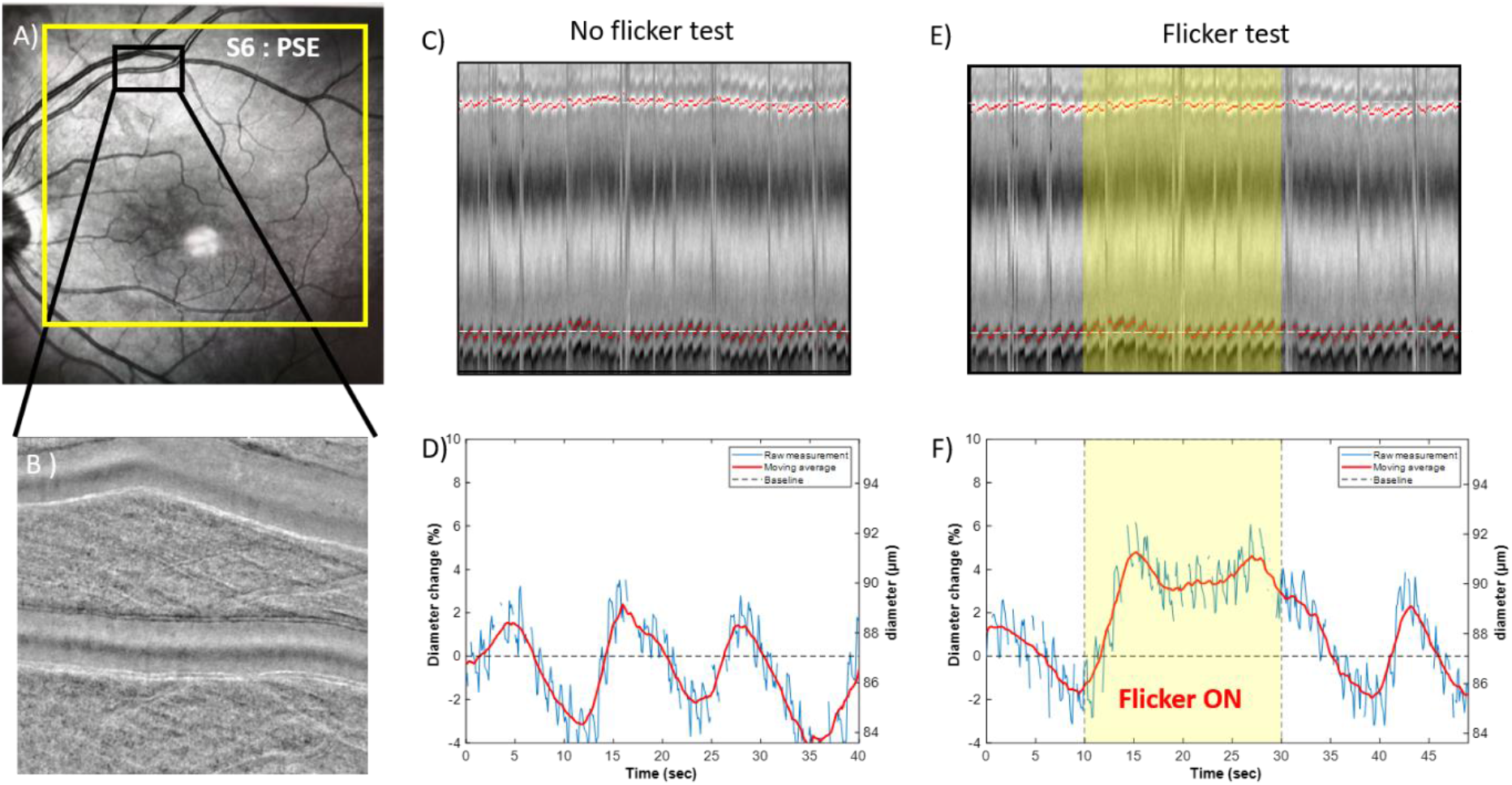

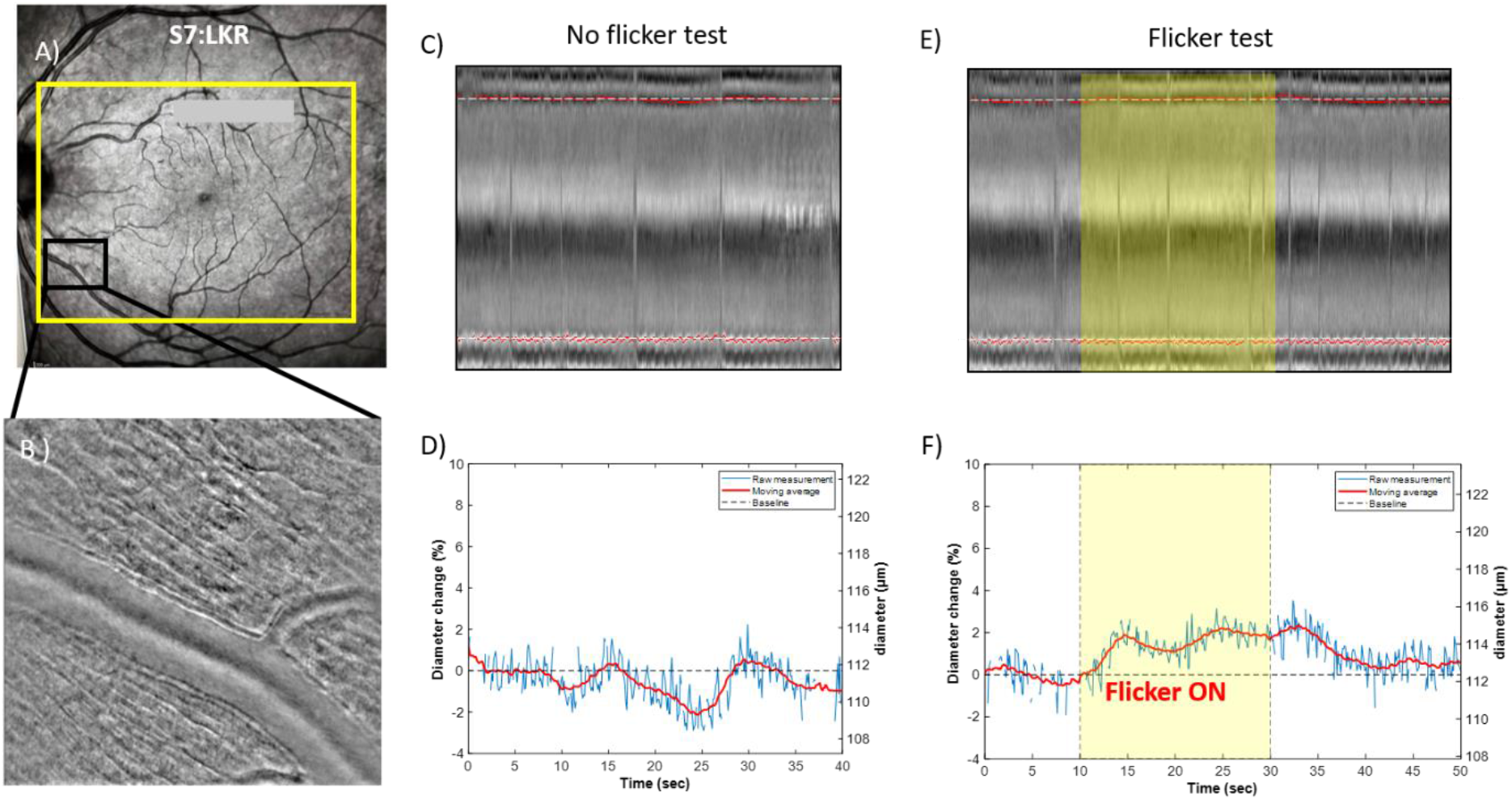

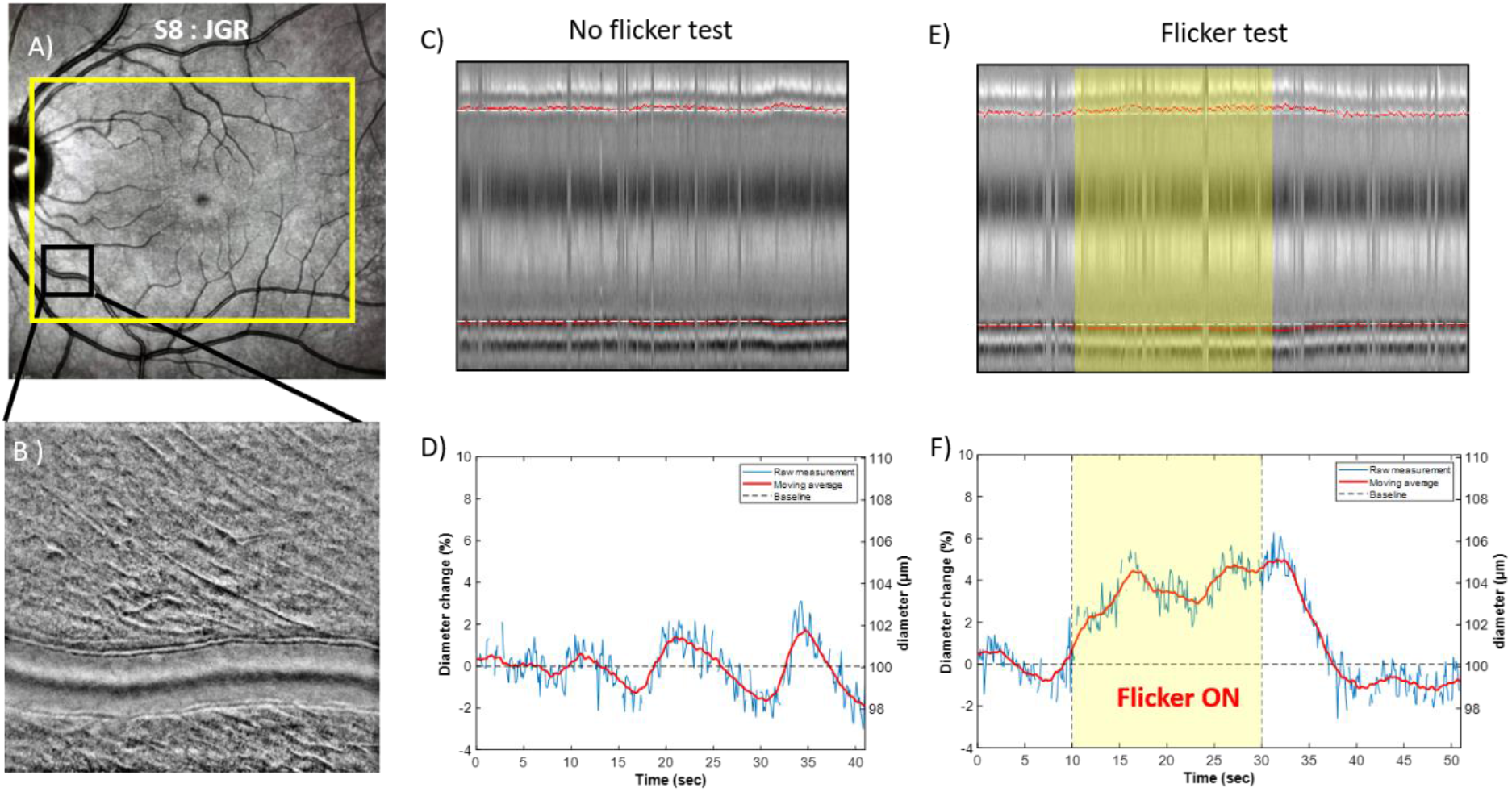

## Notes

### Competing Interest Statement

The authors have declared no competing interest.

